# Genomic Surveillance of Yellow Fever Virus Epizootic in São Paulo, Brazil, 2016 – 2018

**DOI:** 10.1101/645341

**Authors:** S. C. Hill, R. P. de Souza, J. Thézé, I. Claro, R. S. Aguiar, L. Abade, F. C. P. Santos, M. S. Cunha, J. S. Nogueira, F. C. S. Salles, I. M. Rocco, A. Y. Maeda, F. G. S. Vasami, L. du Plessis, P. P. Silveira, J. de Goes, J. Quick, N. C. C. A. Fernandes, J. M. Guerra, R. A. Réssio, M. Giovanetti, L. C. J. Alcantara, C. S. Cirqueira, J.D. Delgado, F. L. L. Macedo, M. C. S. T. Timenetsky, R. de Paula, R. Spinola, J.T. Telles de Deus, L.F. Mucci, R.M. Tubaki, R.M.T. Menezes, P.L. Ramos, A. L. Abreu, L. N. Cruz, N. Loman, S. Dellicour, O. G. Pybus, E. C. Sabino, N. R. Faria

## Abstract

São Paulo (SP), a densely inhabited state in southeast Brazil that contains the fourth most populated city in the world, recently experienced its largest yellow fever virus (YFV) outbreak in decades. YFV does not normally circulate extensively in SP, so most people were unvaccinated when the outbreak began. Surveillance in non-human primates (NHPs) is important for determining the magnitude and geographic extent of an epizootic, thereby helping to evaluate the risk of YFV spillover to humans. Data from infected NHPs can give more accurate insights into YFV spread than when using data from human cases alone. To contextualise human cases, identify epizootic foci and uncover the rate and direction of YFV spread in SP, we generated and analysed virus genomic data and epizootic case data from NHP in SP. We report the occurrence of three spatiotemporally distinct phases of the outbreak in SP prior to February 2018. We generated 51 new virus genomes from YFV positive cases identified in 23 different municipalities in SP, mostly sampled from non-human primates between October 2016 and January 2018. Although we observe substantial heterogeneity in lineage dispersal velocities between phylogenetic branches, continuous phylogeographic analyses of generated YFV genomes suggest that YFV lineages spread in São Paulo state at a mean rate of approximately 1km per day during all phases of the outbreak. Viral lineages from the first epizootic phase in northern São Paulo subsequently dispersed towards the south of the state to cause the second and third epizootic phases there. This alters our understanding of how YFV was introduced into the densely populated south of SP state. Our results shed light on the sylvatic transmission of yellow fever in highly fragmented forested regions in SP state and highlight the importance of continued surveillance of zoonotic pathogens in sentinel species.

**Author’s Summary:** Since July 2016, the southeast region of Brazil has experienced the largest yellow fever virus (YFV) outbreak in decades. São Paulo is the most densely populated state in southeast Brazil. The outbreak has caused serious public health concern in the state, as YFV does not normally circulate widely there and most of the 21 million inhabitants were correspondingly unvaccinated against YFV when the outbreak began. In Brazil, YFV typically circulates among non-human primates, and human cases represent isolated spillover events from this predominantly sylvatic cycle. Understanding the epidemiological dynamics and spread of YFV in non-human primates is therefore critical for contextualising human cases, and guiding vaccination strategies that can better protect local human populations. Here, we aim to contextualise human cases, identify epizootic foci and uncover the rate and direction of YFV spread in SP. We analyse the geographic and temporal distribution of observed cases of YFV in non-human primates in São Paulo state, and identify three distinct phases of the epizootic. We generate sequence data from 51 YFV-positive cases and perform phylogenetic and phylogeographic analyses aimed at understanding the spatial spread of YFV in São Paulo state. Analyses of these data indicate that YFV spread from the north of São Paulo state into more densely populated southern regions. Although we observe substantial heterogeneity in the rate at which different sampled YFV lineages spread, the typical rate of spread was low with a mean rate of ~1 km per day. This is consistent with a scenario in which the majority of transmission events occurred between non-human primates and sylvatic vectors across forested patches.

**Article Summary Line:** Genomic surveillance of yellow fever in São Paulo during the 2016-2018 epizootic

## Introduction

Yellow fever (YF) is an acute hemorrhagic disease caused by the yellow fever virus (YFV), a single-stranded positive-sense RNA virus from the *Flavivirus* genus. Yellow fever is endemic to the American and African tropics, where nearly 400 million people are estimated to be at risk of infection (1). Clinical manifestations of YF in humans range from inapparent or mild disease in up to 80% of infected cases, to severe hepatitis and hemorrhagic disease. The fatality rate among patients who develop visceral disease can range from 20% to 60% (2). YF is preventable in humans by administration of a single dose of an extremely effective vaccine that provides life-long protection against the disease.

YFV is primarily transmitted in the Americas in a sylvatic cycle between non-human primates and tree-dwelling mosquitoes (primarily *Haemagogus janthinomys, Haemagogus leucocelaenus* and more rarely mosquitoes of the *Sabethes* genus) (3–5). The urban cycle of YFV, in which YFV is transmitted between humans and *Aedes aegypti* mosquitoes, has not been observed in the Americas since the 1940s (2,6). In Brazil, outbreaks typically initiate amongst NHPs in neotropical forests and cause subsequent human epidemics that mostly affect unvaccinated male adults in rural areas. YFV infection tends to be detected earlier in NHPs before occasional human spillover infections are observed, and infection location is typically more precisely ascertained for NHPs than for humans. Surveillance of primate epizootics is therefore critical to alert us to the possibility of future human infections (7), and to provide the clearest possible indicator of spatiotemporal disease spread.

From July 2016, the southeast region of Brazil suffered the largest YFV outbreak observed in the Americas in decades. The epizootic/epidemic was caused by a rapidly-spreading lineage of the South American 1 (SA1) genotype that is thought to have originated in the Amazon basin (8). Between December 2016 and June 2019, the Brazilian Ministry of Health confirmed at least 2251 human YFV cases and 1567 epizootic events in non-human primates across Brazil (9). São Paulo state is a densely populated state of southeast Brazil that contains one of the world’s largest urban conurbations. Available reports from São Paulo suggest that the state health authorities confirmed around 875 cases of YFV in NHPs between July 2016 and 18^th^ November 2019 and 624 cases of YFV in humans between January 2017 and 18^th^ November 2019 (10–12). YFV genomes from 36 human cases sampled during the outbreak have recently been reported from SP (13). However, despite the magnitude and expansion of the epizootic in São Paulo (SP) state, remarkably little is known about the origins, spread or genetic diversity of YFV in NHP populations in the region.

Analytical insights into the spread of YFV among NHP populations can guide public health efforts more effectively than those based on human cases alone. This is because routes and rate of YFV spread estimated solely from human cases can be biased by substantial human travel away from the location of infection, or obscured by the lack of human cases in well-vaccinated regions where epizootics are nevertheless ongoing. Epizootic foci can be used to determine which human populations may be most urgently in need of vaccination. Understanding how YFV is introduced into new areas, particularly highly urbanised areas such as São Paulo city, is important for designing strategies that can interrupt these introductions. Estimates of the rate of YFV spread can help public health organisations ensure healthcare preparedness is achieved before the anticipated onset of local human cases. To address these questions, we report epizoological case data from NHP populations in São Paulo state from January 2016 - February 2018. We sequence YFV from NHPs in São Paulo state using a previously-developed portable sequencing approach. We analyse case data and conducted virus phylogeographic analyses to characterize the spread of YFV in São Paulo state during the first waves of the outbreak.

## Methods

### Sample collection and mapping of non-human primate cases

Notifications of dead NHPs were made to the São Paulo’s State Epidemiological Health Department through the Information System for Notifiable Diseases (SINAN). Quantum GIS (14) was used to create a choropleth map depicting the geographic distribution and number of confirmed cases across different municipalities in São Paulo state. The number of reported NHPs varied over time in both northern and southern parts of the state (**Fig. S1**).

### Identification of YFV positive cases

Reported NHP carcasses were tested for YFV. Not all reported carcasses were subsequently tested for YFV and therefore the number of positive animals reported here likely underestimates YFV incidence in NHPs in São Paulo. The proportion of all cases in our data changed over time, such that proportionally fewer reported NHPs were tested in late 2017 and early 2018 than at an early stage of outbreak emergence (**Fig. S1**). Despite this, the proportion of all carcasses that were tested following reporting was relatively consistent across each municipality (**Fig. S2**).

YFV was confirmed by a positive result in at least one of the three methods described below; immunohistochemistry, immunofluorescence, reverse transcription quantitative PCR (RT-qPCR), or epidemiological linkage.

### Histopathology and immunohistochemistry for NHP cases

All samples were subjected to histopathology and immunohistochemistry examination. Samples of brain, heart, lung, liver, spleen and kidney from NHPs were fixed in formaldehyde and embedded in paraffin. Histological sections of these tissues were stained with hematoxylin and eosin and examined on a microscope. Indirect immunohistochemistry was used to detect the presence of the yellow fever virus antigen. Sections of liver (0.3uM) were placed on slides coated with silane and were treated with an in-house polyclonal anti-YFV antibody that was produced in mice and used at 1/30,000 dilution. The slides were treated with anti-mouse secondary antibodies, linked to either horseradish peroxidase (Reveal HRP Spring polymer System, Spring) or to alkaline phosphate (Link MACH4 universal AP Polymer and Polymer MACH4 universal AP, Biocare). Chromogenic detection of the presence of YFV was subsequently conducted using the substrates 3,3’-diaminobenzidine or Fast Red (Warp Red, Biocare), respectively.

### Indirect immunofluorescence

Samples of NHP blood or serum, and tissue material suspensions obtained from autopsies, were tested using a standardized indirect immunofluorescence technique (15). An in-house polyclonal anti-flavivirus antibody (anti-DENV 1-4) and an anti-mouse IgG-FITC antibody (Sigma) were used. Positive samples were typed by indirect immunofluorescence with monoclonal antibodies for YFV (Biomanguinhos, Rio de Janeiro).

### RT-qPCR

Total RNA was extracted from tissue and serum samples using two commercial kits according to the manufacturer’s instructions: QIAamp^®^ RNA Blood for tissues and QIAamp^®^ Viral RNA Kit for serum (Qiagen Inc., Germany). Viral RNA was detected using two previously published RT-qPCR techniques (16,17).

### Epidemiological linkage

Where multiple NHPs from the same species were found dead at the same time and location, the number of carcasses was recorded but typically only one or a few animals were tested. In these instances, untested animals were on rare occasions considered as confirmed cases on the basis of their spatiotemporal association with with tested, confirmed cases.

### Sample collection of human cases

We randomly selected 5 samples from confirmed human cases collected in municipalities of São Paulo state (**Table S1**) that had been tested using the RT-qPCR methods, above. These samples were collected in several municipalities of São Paulo state and sent for molecular diagnostics at Instituto Adolfo Lutz.

### Ethics statement

Research on human cases was supported by the Brazilian Ministry of Health (MoH) as part of arboviral genomic surveillance efforts within the terms of Resolution 510/2016 of CONEP (Comissão Nacional de Ética em Pesquisa, Ministério da Saúde; National Ethical Committee for Research, Ministry of Health). Only naturally deceased NHPs were sampled for YFV surveillance. The surveillance protocol for dead NHPs was approved by the Ethics Committee for the use of Animals in Research, Instituto Adolfo Lutz, under the numbers 0135D/2012 and 020G/2014.

### MinION genome sequencing

A selection of positive samples (**Fig. S3** and **Table S1**) was sequenced using a rapid whole-genome sequencing protocol that has been previously validated and successfully applied in Brazil (8,18). NHPs samples were selected to have RT-qPCR CT values <25 to facilitate genomic amplification and sequencing. Because few samples were available, human samples were randomly selected from positive cases with CT <37. In brief, cDNA was produced from viral RNA using random hexamers and the Protoscript II First Strand cDNA synthesis kit (NEB). The genome was amplified using a multiplex PCR scheme designed to produce overlapping 500bp amplicons across the whole coding region of the recent South American genotype I outbreak clade (8,18). PCR products were quantified, barcoded using the Oxford Nanopore Technologies (ONT) Native Barcoding Kit (NBD103, or NBD103 and NBD114, depending on sequencing date), and pooled in an equimolar fashion. Sequencing libraries consisting of 10-24 samples per library were constructed using a Ligation Sequencing kit (ONT, SQK-LSK108 or SQK-LSK109, depending on sequencing date). Sequencing was performed on a ONT FLO-MIN106 flow cell for up to 48h as described previously (8).

### Generation of consensus sequences

Previously published approaches were used to produce YFV consensus sequences (18,19). In brief, raw files were basecalled using Albacore or Guppy version 3.0.3 (Oxford Nanopore Technologies). Reads were demultiplexed and trimmed of adaptor and barcode sequences using Porechop (version 0.2.3, https://github.com.rrwick/Porechop). Trimmed reads for each barcoded sample were mapped to a reference genome (accession number JF912190) using *bwa* (20), and primer binding sites were trimmed from reads. Single nucleotide variants to the reference genome were detected using nanopolish variant calling (version 0.11.1) (21). Consensus sequences were generated using identified variants. Sites with <20X coverage were replaced with ambiguity code N. Sequencing statistics can be found in **Table S2**. Accession numbers of newly generated sequences can be found in **Table S1**.

### Phylogenetic analyses

The genome sequences generated here *(n* = 51) were combined with all previously published SA1 genotype YFV genomes available to 31 December 2019 (n = 173). This dataset was curated to remove all sequences shorter than 70% of the whole coding sequence length, those with missing metadata, and those with abnormal genomic sequences typical of sequencing or assembly errors. The full dataset analysed here therefore contains 224 YFV sequences, as shown in **Fig. S4**. Sequences were aligned using MAFFT v.7 (22). The best-fit substitution model was selected using ModelTest-NG (23). Maximum likelihood (ML) phylogenetic trees were estimated using RAxML v.8 (24) under a GTR + I + Γ_4_ nucleotide substitution model. Statistical support for phylogenetic nodes was estimated using a ML bootstrap approach with 100 replicates.

### Phylogeographic analyses

To investigate the spread of YFV in SP, we analysed the SP outbreak clade in more detail. We focus on two datasets; the ‘full’ SP outbreak clade, represented by 99 sequences sampled from 2016-2018 (shown as individual taxa in a ML phylogeny in **Fig. 3)**, and a ‘geographically restricted’ dataset in which 4 sequences are removed that appear to be clear spatial exports from the main SP epizootic to distant locations in Goiás, Espírito Santo and Minas Gerais, with little phylogenetic evidence of extensive local transmission (**Fig. 4**). These spatial outliers were removed so that rare, long-distance movements with little or no observed onwards transmission did not impact estimates of branch dispersal rate or maximal wavefront distance.

Geo-referenced and time-stamped sequences were analysed in BEAST v. 1.10.4 (25) using the BEAGLE library to enhance computational speed (26). The uncorrelated lognormally distributed relaxed clock model (27), a skygrid coalescent tree prior (28) and a continuous phylogeographic model that uses a relaxed random walk (RRW) to model the spatial diffusion of lineages were used. Contrary to a standard Brownian diffusion model that is based on a strong assumption of homogeneous dispersal velocity in all phylogenetic branches, the RRW model is a flexible model that specifically allows inference of heterogeneous, branch-specific dispersal velocities. Dispersal velocity variation among lineages was here represented using a Cauchy distribution (29,30). The default settings were used for the diffusion rate. Uniform priors were used to allow date uncertainty for those sequences where exact sampling day was unknown, and a jitter added to tips with non-unique co-ordinates (window size of 0.2 decimal degrees).

Virus diffusion through time and space was summarised using 1000 phylogenies sampled at regular intervals from the posterior distribution (after exclusion of 10% burn-in). The R package “seraphim” (31) was used to extract the spatiotemporal information contained in these phylogenies, whose branches can be considered as movement vectors (each having start and end spatial coordinates, and start and end dates, in decimal units). The package was also used to estimate statistics of spatial dissemination, such as mean branch dispersal velocity (**Fig. 6**), change in the maximal wavefront distance from epizootic origin, and an estimate of subsequent wavefront dispersal velocity (**Fig. 6**) (31). The maximum wavefront distance is the distance between the epidemic origin (here approximated as the inferred tree root location) and the furthest location at which virus transmission has ever occurred. Wavefront velocity is the rate at which the maximum wavefront distance increases over time. In contrast, branch dispersal velocity captures the rate of movement of any lineage, regardless of whether or not it expands the epidemic wavefront. The results shown in **Fig. 4** were generated using the “spreadGraphic2” R function (31).

### Data availability

Epidemiological data, genomic data and XML files analysed in this study are available in the GitHub repository (https://github.com/CADDE-CENTRE). New sequences have been deposited in GenBank under accession numbers MH030049-MH030090, MH193173-MH193175 and MT497520-MMT497525.

## Results

Liver, brain or blood samples from 591 NHPs were considered positive for YFV between epidemiological week 29 of 2016 to week 4 of 2018. Samples were confirmed as YFV cases using immunohistochemistry (*n* = 268), RT-qPCR (*n* = 113), both immunohistochemistry and RT-qPCR (n = 198) or in rare instances by epidemiological linkage to another positive NHP (n = 12). Most confirmed YFV-positive NHP cases were from animals of the *Alouatta* genus (88%; 403 of 459 cases for which genus information was available), followed by *Callithrix* (8%; 35/459), *Callicebus* (2%; 9/459), *Cebus* (2%; 9/459) and *Sapajus* (0.7%; 3/459).

Our results reveal three main epizootic phases, including a primary phase of low reported NHP case counts in northern SP state, followed by two phases where higher numbers of NHP cases were reported almost exclusively in the south the of state. Phase 1 runs from July 2016 to Jan 2017 (6%, 33/591 cases). Phase 2 runs from February to July 2017 (20%, 119/591 cases) and phase 3 from July 2017 to at least February 2018 (74%, 439/591 cases) (**Fig. 1A**). These transition dates were chosen to capture the spatial shift between phase 1 and phase 2 (**Fig. 1B**), and to occur centrally within the low case count period that occurs between phase 2 and 3 (**Fig. 1A**), but the exact dates adopted here are somewhat arbitrary. The peaks of epizootic phases 2 and 3 were mid-April 2017 and late November 2017, respectively. Proportionally fewer NHP carcasses were tested in phase 3 than in phase 1 or 2, and in January and February 2018 than in many earlier months (**Fig. S1**). It is therefore possible that phase 3 may have been of longer duration and proportionally greater magnitude than we report here.

**Figure 1.**
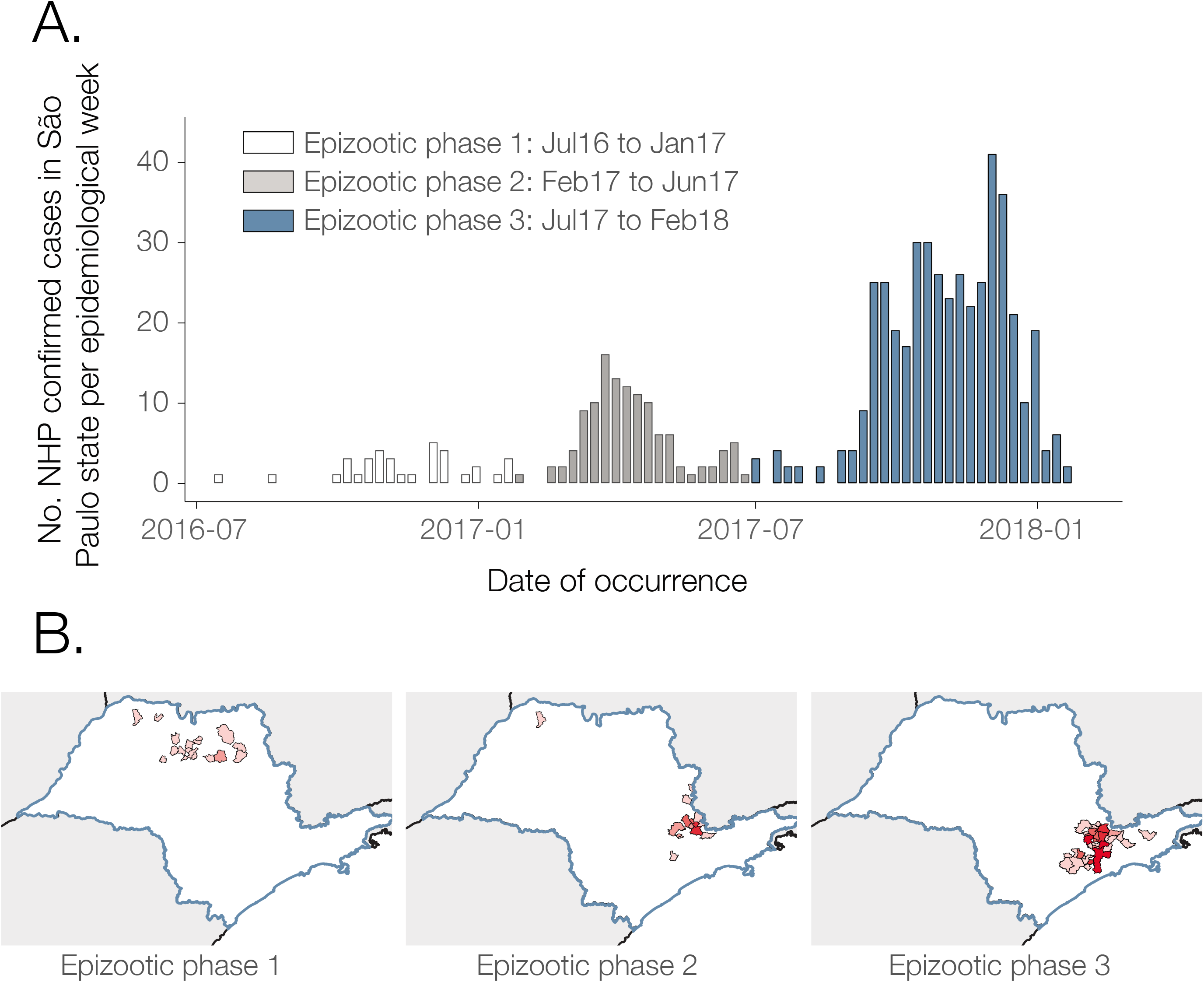
Distribution of weekly YFV cases. **A)**. NHP YFV cases diagnosed by RT-qPCR per week. **B)**. Choropleth map of SP municipalities reporting positive NHP YFV cases during each outbreak phase. Samples SA131 and Y37-244 are indicated.

NHP cases were geographically widespread across 57 municipalities of SP to February 2018, with most reported cases concentrated around the southeast region of the state (**Fig. 2**). The greatest number of confirmed cases were observed in the municipalities of Mairiporã (*n* = 88), Jundiaí (*n* = 68), São Paulo (*n* = 62), Bragança Paulista (*n* = 48), and Atibaia (*n* = 36). There is a clear distinction between the geographic location of cases in epizootic phase 1, and those in phases 2 and 3 (**Fig. 1B**). Specifically, almost all cases that form part of phase 1 occur in a geographically distinct cluster of municipalities in the north of São Paulo state (**Fig. 1B**; **Fig. 2)**. On the other hand, nearly all cases of YFV detected after the first phase occurred in municipalities in the South of São Paulo state (within 200km of São Paulo city. (**Fig. 1B and Fig. 2**).

**Figure 2.**
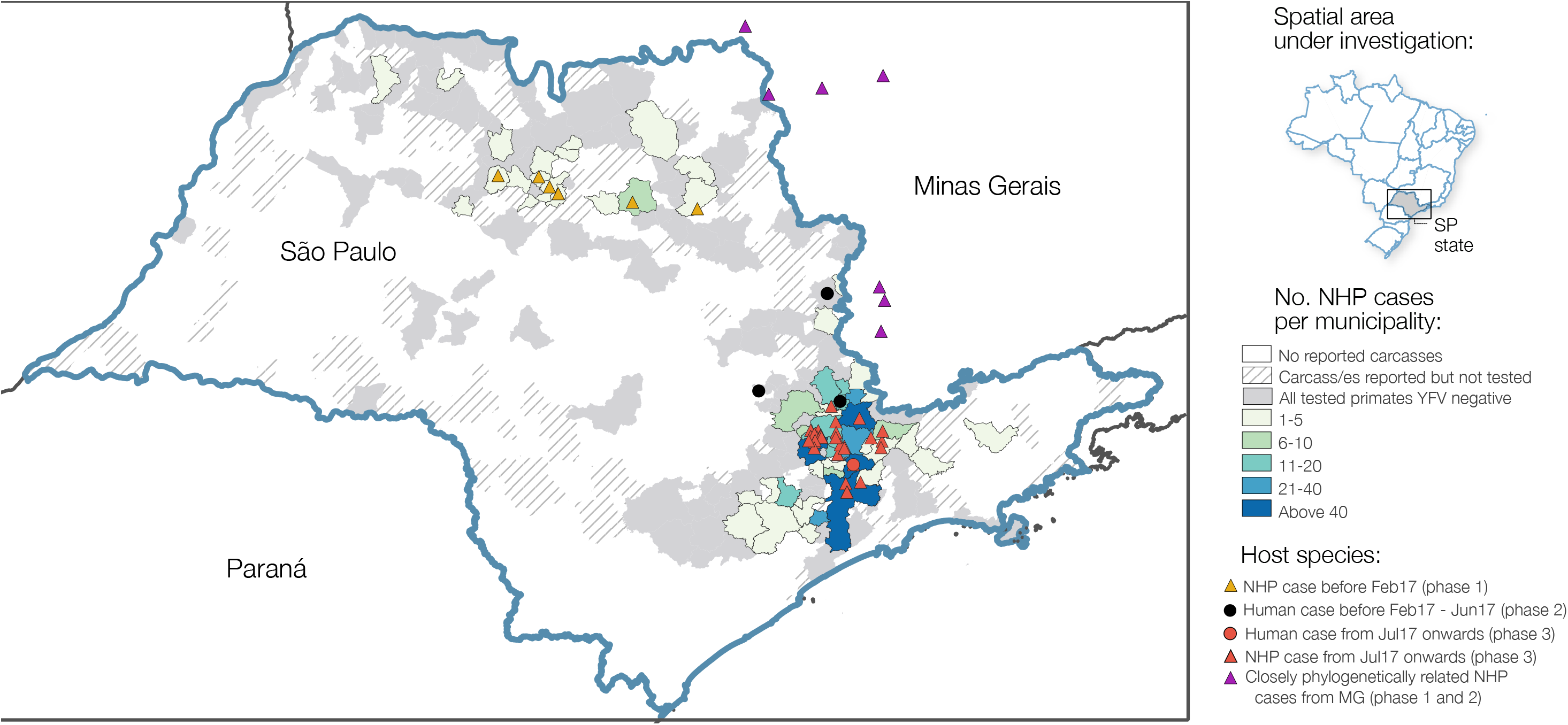
Choropleth map of the distribution of confirmed NHP cases per municipality in São Paulo state between July 2016 and February 2018. Circles depict locations of those cases from humans that were sequenced as part of this study; yellow and red triangles depict locations in São Paulo state from which non-human primate genome sequences have been generated as part of this study. Eight samples from 7 municipalities in MG that form a clade with our sequences from SP and that were sequenced by previous studies (8,42) are shown in purple triangles.

To investigate the source and transmission of YFV during all three epizootic waves, and to characterise the genetic diversity of the virus circulating in NHP populations across SP state, we generated whole YFV genomes for 46 RT-qPCR positive NHP samples (median Ct-values of 14, range 9-25) from 20 municipalities using a previously described MinION sequencing protocol (8,18). We also sequenced five whole genomes from human samples collected in four municipalities of São Paulo state (median Ct-values of 34, range 32-37). In total, this represents genomes from 51 samples collected across 23 municipalities between October 2016 and January 2018 (**Fig. S3)**.

We first used maximum likelihood methods to estimate phylogenetic relationships amongst all 224 sequences available from SA1 in South America. Our genetic analyses indicate a strong phylogenetic spatial structure of the ongoing epizootic in SP state, with 43 of 46 sequences from NHPs in SP clustering in a single clade (bootstrap score = 98%), previously named as lineage YFVMG/SP (32). This clade contains sequences sampled in SP state, and an additional 8 sequences sampled in Minas Gerais that were all from within 100km of SP state (**Fig. 3**, see **Fig. 2** for locations of close municipalities and **Fig. S4** for the full phylogeny containing all sequences).

**Figure 3.**
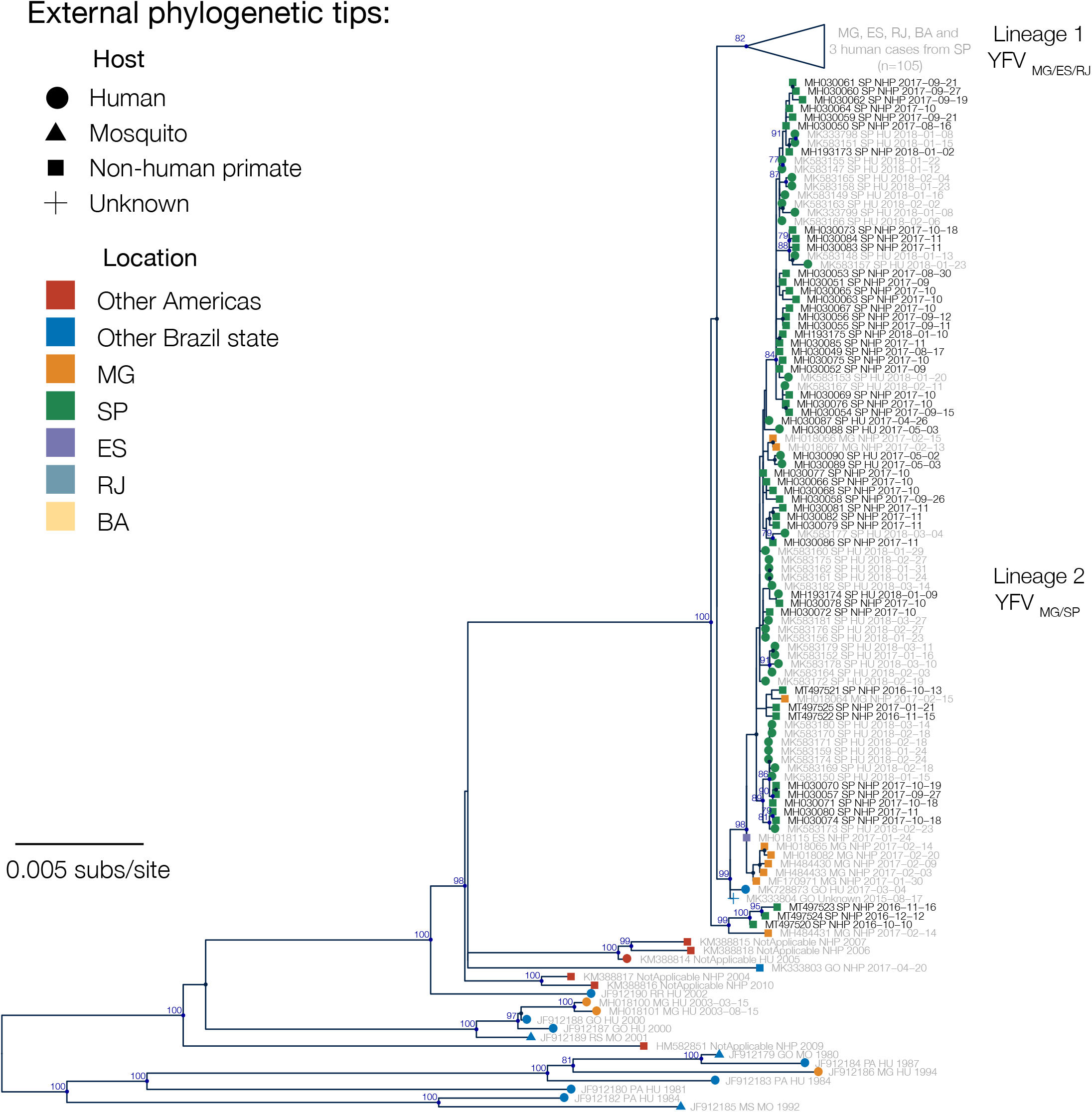
Maximum likelihood phylogenetic tree of YFV in São Paulo, Brazil. Nodes with bootstrap support (>50%) are coloured dark blue, and bootstrap scores are provided for highly supported nodes (> = 75%). Sequences generated in this study have black tip labels (n = 51). One clade (previously denoted lineage YFVMG/ES/RJ (32)) is collapsed for clarity. This clade contains three phylogenetically unlinked sequences that were sampled from humans in SP, but for all three cases patient travel histories indicate that YFV was acquired elsewhere. Samples SA131 and Y37-244 are indicated with asterisks and discussed in the Discussion.

Sequences from 6 NHPs sampled during the first epizootic phase (October to December 2016) form two separate phylogenetic clades in the maximum likelihood tree, one of which appears basal to the 2016-2019 outbreak clade in southeast Brazil. Whilst bootstrap support for this structure is relatively poor (bootstrap score =67%), its presence raises the possibility that two distinct YFV lineages may have co-circulated in northern São Paulo state during the first low-intensity epizootic phase in late 2016.

We used a continuous diffusion model to investigate the spatiotemporal spread of YFV. Our analyses were unable to accurately determine the route via which YFV was introduced to SP state.

Although the root of the MCC tree is placed in northwestern MG (red dot, **Fig. 4**), the spatial uncertainty in this root location is very high. In addition, posterior support for phylogenetic grouping of sequences from Goiás state with those from northern São Paulo is very weak (0.12), and as such this reconstructed lineage movement should be interpreted with caution. Whilst we were unable to identify the route of introduction to northern SP, our analyses are better able to resolve subsequent viral dispersal trajectories from the north to the south of the state. Specifically, YFV appears to have disseminated from northern SP into geographically neighbouring areas of western MG and into the south of the state.

Both those analyses including and discarding spatially-outlying sequences (see **Fig. 4** for locations of the discarded sequences) suggest that the mean branch velocity was ~1 km/day (geographically restricted dataset: 0.83 km/day, 95% highest posterior density (HPD), 0.50-1.53, full dataset: 0.95 km/day, 95% HPD 0.53-2.42). The mean velocity did not vary significantly between the three epizootic phases observed in SP (**Table S3**). There is, however, heterogeneity in the dispersal velocity of different branches, such that most phylogenetic branches have lineage dispersal velocities that are somewhat lower than the mean, and a few are associated with rapid dispersal (**Fig. 5**).

**Figure 4.**
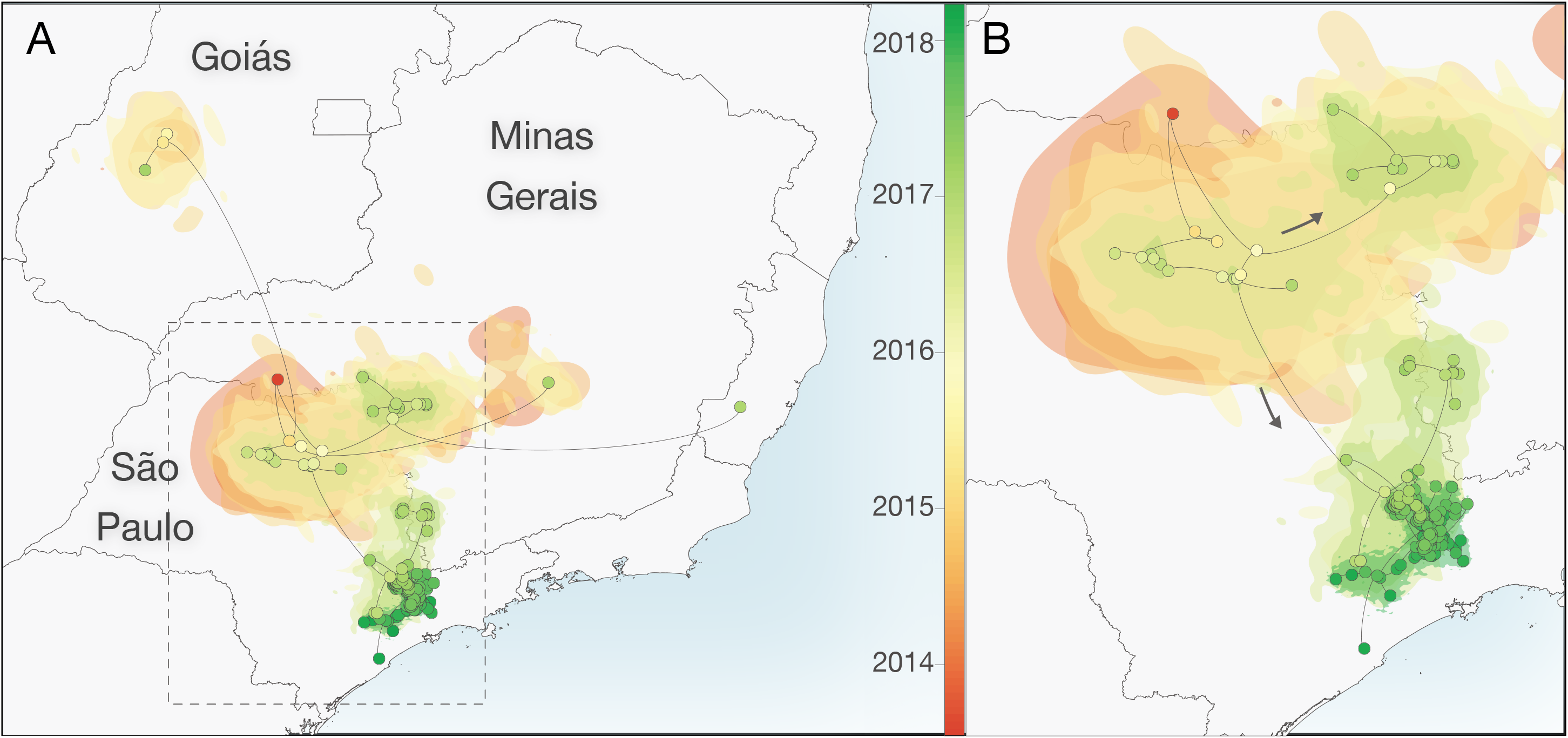
Reconstructed spatiotemporal diffusion of YFV in São Paulo. Phylogenetic branches are mapped in space according to the location of phylogenetic nodes and tips (circles). Shaded regions coloured according to time show 80% highest posterior density contours calculated using bivariate kernel density estimation of the location of all nodes observed within defined 6-month time intervals in a subset of 1000 trees. Illustrated nodes are from the MCC tree and are coloured according to time. **A)** Spatiotemporal diffusion of taxa based on analysis of the full dataset (n = 99 sequences). **B)** Expansion of region highlighted in **A** without the long-distance outlier sequences.

**Figure 5.**
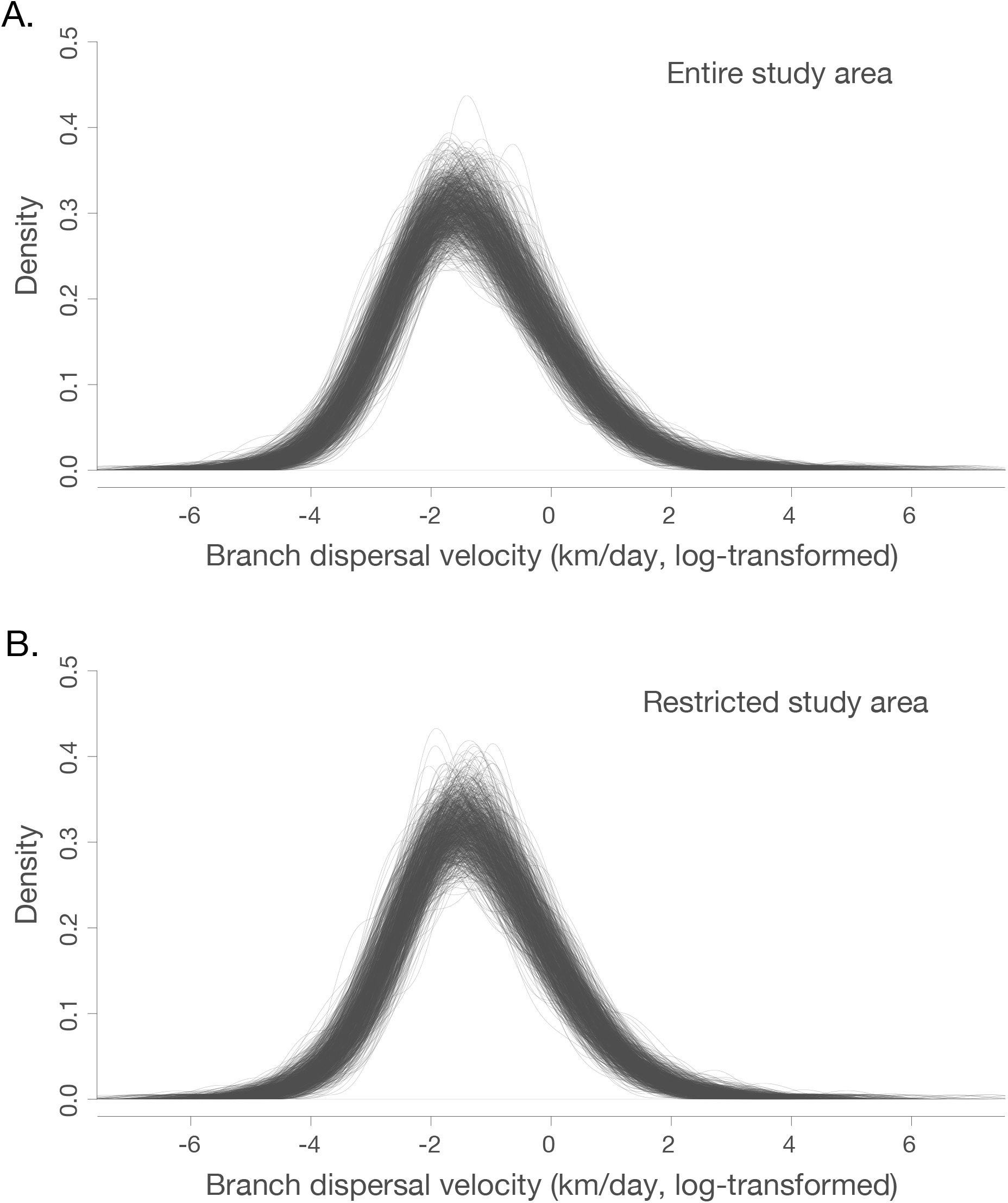
Density of branch dispersal velocity in entire (panel A) and restricted (panel B) study area. Branch dispersal velocities are natural log transformed. Each individual trace represents branch dispersal velocity density of one tree from the generated posterior distribution of phylogenetic trees.

Considering all sequences, the wavefront velocity varies across time **(Fig. 6**). However, rare distant outlier sequences can heavily inform the maximum wavefront distance, even if associated expansive virus lineages do no contribute substantially to the overall epizootic and only represent isolated cases in sampled outlier locations. Considering sequences in the spatially restricted area only (see **Fig. 4** for locations), the posterior intervals on the wavefront velocity are relatively wide and we therefore cannot exclude that the wavefront velocity of the majority of the sampled epizootic is homogeneous across time (**Fig. 6**). In this dataset, the epidemic wavefront moved on average only 0.4 km/day (95% HPD, 0.30-0.53) during January 2015 to January 2017 (**Fig. 6**).

**Figure 6.**
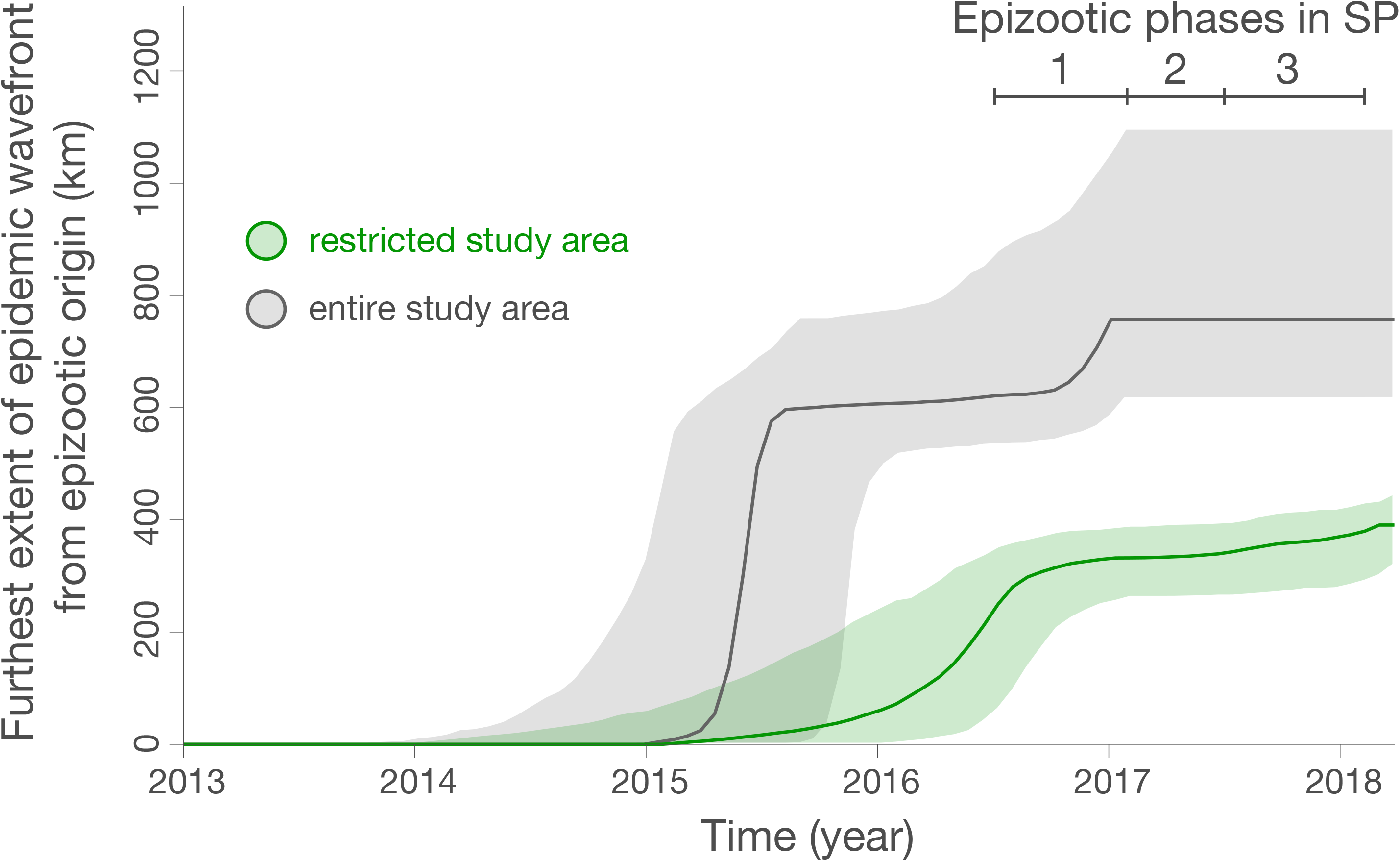
Spatial wavefront distance from epidemic origin over time for all studied sequences (grey), and those in the restricted area shown in **Fig. 4B** (green), as estimated with the “seraphim” package. The identified epizootic phases are indicated by bars in the upper right. Ribbons show 95% HPDs.

## Discussion

In this study we analyse data on the epizootic intensity and spatiotemporal distribution of YFV cases in NHPs in São Paulo state, Brazil, during 2016-2018. Our results demonstrate the existence of three distinct phases of the epizootic to early 2018. Specifically, we observe an initial phase in which a small number of cases were identified in the northern region of SP state during late 2016. This period was followed by two larger epizootic phases; the first from February 2017 — July 2017, and the second from July 2017 to at least February 2018 (the date of the most recent data available for this study). During the second and third phases, a large number of cases were observed in the south of SP state. We generated 51 novel YFV genomes from samples collected in 23 different municipalities in São Paulo state. The majority of sequenced viruses were sampled from NHPs of the genus *Alouatta* from late 2016 to early 2018, and represent the first and third epizootic phases identified here (**Fig. S3, Fig. 1**). Phylogeographic analyses of these data indicate that after being introduced to the north of SP state, YFV lineages spread southwards and initiated epizootics in the south of the state. Lineages spread at a mean branch velocity of approximately 1km per day (**Fig. 4 and Fig. 5**), though there is substantial heterogeneity in branch dispersal velocity (**Fig. 5**).

Previous analyses have hypothesised that the epizootic in the densely populated south of SP originated in Minas Gerais (13), but those analyses were performed without any data originating from the north of SP state. Here, we revise this hypothesis and suggest that the epizootic in the densely populated south of SP (epizootic phase 2 and 3) may instead have originated in the north of the state. However, we note that few YFV-positive carcasses were identified in the centre-east of SP (**Fig. 2**) and genomes sampled from NHP in the north and south of SP are connected by a single branch in the phylogenetic tree, with no genomes sampled in between. As such, it is impossible to identify the exact route or mode of YFV spread through the state. Several possibilities might explain the movement of YFV between northern and southern SP: (i), YFV-positive NHPs were present but were not identified in a corridor in the eastern SP region due to limited sampling (**Fig. 2**), (ii), YFV moved via an as-yet-unsampled ‘stepping-stone’ NHP populations in neighbouring MG, or (iii), YFV was directly transmitted between the two locations through human activities, without the infection of intermediate NHP populations. The northern regions of SP that reported YFV in 2016-2017 are similar to those affected by YFV in 2008 and 2000 (33,34), and it is therefore plausible that YFV will emerge again there in future. Understanding how and why the outbreak spread southwards more extensively from northern SP during the 2016-2019 outbreak than in previous outbreaks could help inform future surveillance and control efforts.

Determining how YFV is introduced into highly urbanised areas, such as São Paulo city, is important for designing strategies that can effectively interrupt such introductions. One of our sequenced samples, SA131 (accession number MH193175), was collected from an *Alouatta* sp. individual at the Parque Estadual das Fontes do Ipiranga (PEFI) (**Fig. 1** and **Fig. 3**). This park forms part of São Paulo city’s zoo, and represents a 53 km fragmented island of Atlantic forest within the urbanised city metropolis. Surprisingly, phylogenetic analyses show that the YFV genome recovered from sample SA131 clusters together with isolate Y37-244 (accession number MH030085) (bootstrap score for ML tree = 68%, 95% posterior support for MCC tree = 1), collected in Piracaia, 75 km from the park, and not with any of the isolates that were detected in the outskirts of SP (**Fig. 1** and **Fig. 3**). This result could be explained by: (i) incomplete sampling of a wave of continuous transmission among NHPs that live between the two locations; (ii) human-mediated transport of YFV infected NHPs (35), (iii) human-mediated transport of mosquitos, or (iv) introduction of the virus by an asymptomatic human visitor, who carried the virus from Piracaia to PEFI. Scenario (i) seems unlikely since PEFI is an isolated forested fragment with limited connectivity to other forested areas; at this stage, scenarios (ii), (iii) and (iv) all remain possible.

Using YFV genetic data, we estimate that virus lineages dispersed at a mean rate of ~1 km per day during the outbreak in SP (95% HPD, 0.50-1.53). We did not observe substantial differences in the mean lineage dispersal between epizootic phases (**Table S3**), which might have indicated different impact of drivers in different epizootic phases or corresponding affected regions. However, our analyses demonstrated substantial heterogeneity between different phylogenetic branches (**Fig. 5**). Most phylogenetic branches show low lineage velocity (<3 km/day) that are consistent with YFV spread by NHPs and mosquitos between contiguous forested patches. Far more rarely, we observe higher velocity lineage movements. Such movements could indicate human transport of infected NHPs or mosquitos, or long-distance movement of infected humans or mosquitos. We also cannot rule out that terminal phylogenetic branches with unusually high lineage dispersal velocity may have been caused by accidental attribution of incorrect metadata to samples, and efforts should be made to perform confirmatory sequencing of outlier samples and sequence YFV from other samples collected in their vicinity. Greater genomic sampling density is required to better investigate the drivers of such rapid YFV lineage movements.

Our mean estimate of the rate of YFV lineage movement is slightly slower than that those previously estimated for the states of Minas Gerais, Espírito Santo and Rio de Janeiro, and for the southeast region as a whole (~4km/day) (8,13). Phylogenetic estimations of rate of spread often depend on sampling range and tend to be biased when rates are estimated for large geographic areas (36). However, the relatively confined area of YFV circulation suggests that relaxed random walk models used here are a good approximation for viral diffusion in the southeast region of Brazil. Additional analyses of larger datasets are necessary to test whether spatial heterogeneity in NHP density, habitat fragmentation, or impact of human activities, may be responsible for the differences observed in the rate of YFV spread among different areas of southeast Brazil.

The NHP case data presented here (**Fig. 1** and **Fig. 2**) suggest that the epizootic was larger in the southern parts of SP than in the north. We hypothesise that this could be related to different NHP or vector species density or distribution in each area. Specifically, the southern part of SP has large, connected forested areas that support a high diversity and density of NHPs (37,38). In contrast, the north of SP is covered by savannah and semi-deciduous forest (38), which may offer more restricted habitat availability for YFV vectors and amplifying NHP species.

Despite this, the magnitude of the YFV epizootic presented in **Figures 1** and **2** is based on epidemiological surveillance alone, and we caution that any bias in reporting or testing of NHP carcasses would also bias these results. Multiple municipalities reported small numbers of NHP carcasses that could not be tested, and/or tested relatively few carcasses in total. It is therefore possible that municipalities that did not report positive cases (**Fig. 1**) nevertheless had local, undetected epizootics. YFV surveillance based on reporting of dead animals may have led to the appearance of a larger YFV epizootic in the south of SP São Paulo, because animals from the *Alouatta* genus that are more likely to die of YFV infection than other genera (39) are more common in the south of SP (37) than the north. The south of SP is more densely human-populated than the north, so we also cannot exclude that dead NHPs would have been noticed and reported more frequently there. This could lead to the appearance that the outbreak was larger in the south. Reporting of NHP carcasses varies over time (**Fig. S1**). It is impossible to determine to what extent time-vary reporting of NHP carcasses was caused by consistent reporting of time-varying NHP deaths, or by changing frequency at which NHP deaths were reported by human observers. However, our key observation that the epizootic in northern SP was largely extinguished after phase 1 is unlikely to have been severely affected by such surveillance biases, as northern locations continue to test NHP carcasses after this time without detecting positive samples (**Fig. 1**, **Fig. S1**). This finding is also consistent with the spatial distribution of YFV cases in humans (40). Greater efforts to sample mosquito vectors or live NHP during epizootic periods would be important to determine whether reliance on reporting of NHP carcasses biases our understanding of YFV spatial distribution.

To facilitate sequencing, we preferentially studied samples showing low CTs during RT-qPCR. Our selected samples were therefore biased towards those obtained from animals with higher viremia, and hence likely towards animals with greater clinical disease and from the *Alouatta* genus. Further, all phylogeographic inference is conditioned upon sampling, such that we can only reconstruct lineage movement amongst those lineages that we sample. The potential sampling biases that we describe above may have resulted in major or minor differences between the actual viral spread and the inferred dispersal history of lineages (41). Analysis of larger YFV genomic datasets incorporating more sequences from *non-Alouatta* genera (including humans with insubstantial travel history), animals with lower viremia, and animals from genomically unsampled locations is critical to determine whether this could have affected our phylogeographic reconstructions of YFV spread in southeast Brazil.

Our study sheds light on the spatial and temporal dynamics of yellow fever in NHP hosts across SP state. A better understanding of the vectors and host species involved in the persistence of the virus, denser genomic sampling of available positive samples, and targeted investigation into phylogenetic branches associated with long-distance or rapid movements, will be important to understand the drivers of yellow fever transmission and anticipate future outbreaks.

## Acknowledgments

We thank the IAL and the USPTM staff, in particular the team from Núcleo de Doenças de Transmissão Vetorial and from the Centro de Patologia. The research was supported by a Medical Research Council and FAPESP CADDE partnership award (MR/S0195/1), Wellcome Trust and Royal Society Sir Henry Dale Fellowship (grant 204311/Z/16/Z), internal HEFCE GCRF grant 005073, John Fell Research Fund Grant 005166, CNPq #400354/2016-0 and FAPESP# 2016/01735-2, and by the Oxford Martin School. SD is supported by the *Fonds National de la Recherche Scientifique* (FNRS, Belgium) and was previously funded by the *Fonds Wetenschappelijk Onderzoek* (FWO, Belgium). This work was also supported by CNPq/MCTI Decit/SCTIE/MoH (440685/2016-8) and CAPES (88887.130716/2016-00).

## Supporting Information Legends

**Technical Appendix.** Contains additional **Fig. S1- Fig. S4, Table S1** – **S3. Fig. S1**. Reporting and testing of NHPs over time in northern and southern SP. **Fig. S2**: Number of carcasses tested and untested for yellow fever virus per municipality. **Fig. S3**: date of sample collection and number of genomes generated from each host genus. **Fig. S4**: Maximum likelihood phylogeny of all publicly available SA1 YFV sequences from Brazil. **Table S1**: details of the YFV genomes generated in this study. **Table S2**: non-human primate yellow fever virus genome sequences from São Paulo, by host genus. **Table S3:** mean branch dispersal velocity estimates.

## Notes

### Competing Interest Statement

The authors have declared no competing interest.

