## Supplementary Appendix for "Genomic Surveillance of Yellow Fever Virus Epizootic in São Paulo, Brazil, 2016 – 2018"

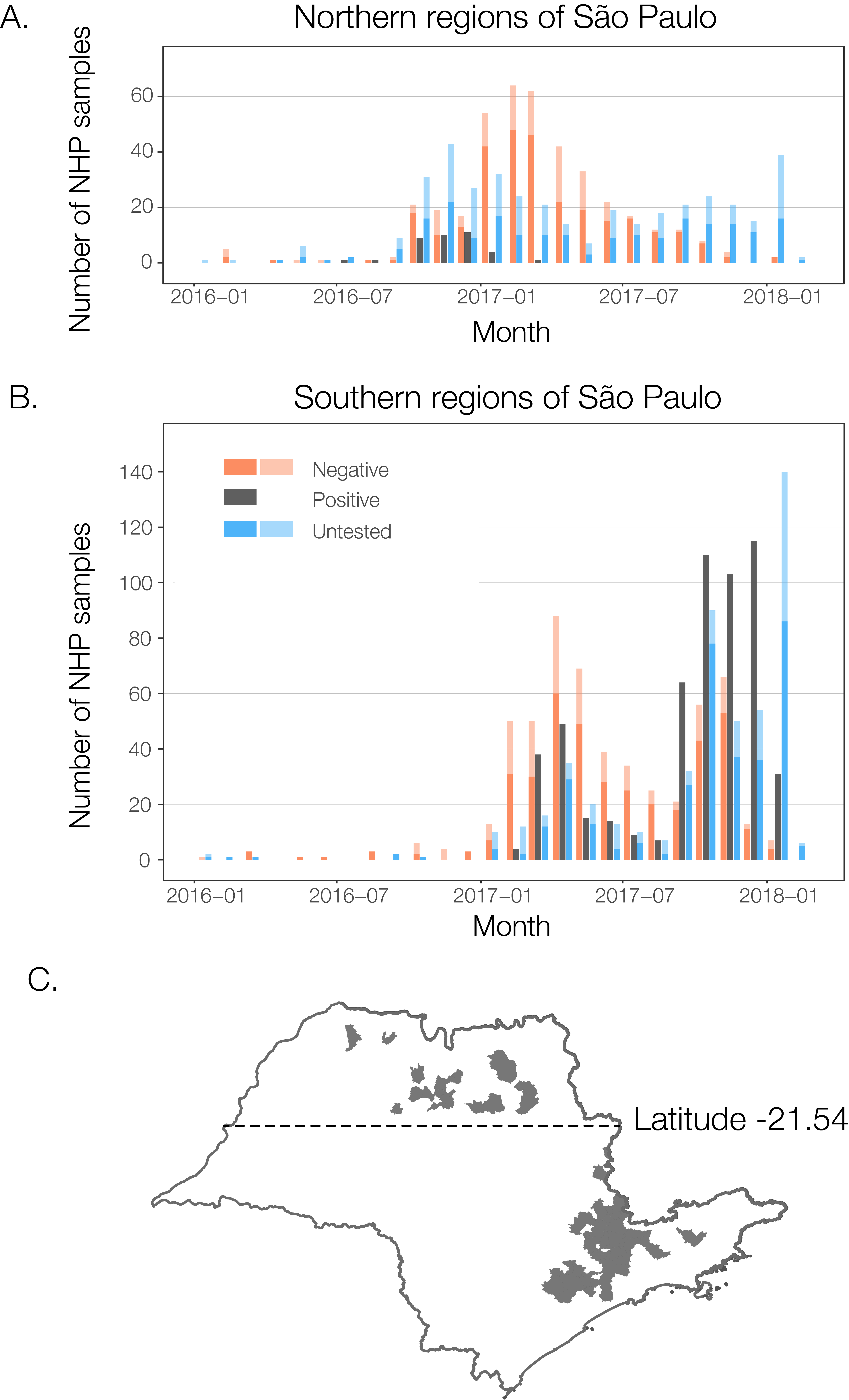

**Figure S1.** Reporting and testing of NHPs over time in northern (**panel A**) and southern (**panel B**) regions of São Paulo state. Here, municipalities where centroids fall north of latitude -21.54 are considered ‘northern’, and others are considered ‘southern’ (latitude indicated on **panel C**, along with locations of municipalities with positive cases in grey). This latitude was chosen to discriminate between locations most affected during phase 1, and those most affected during phase 2 and 3, and is therefore not strictly central in SP. Colours indicate results of testing. Darker shades of each colour in each bar represent results in testing in any municipality that detected positive NHPs during any month, and lighter shades represent cases from those municipalities that never detected positive cases during the displayed period.

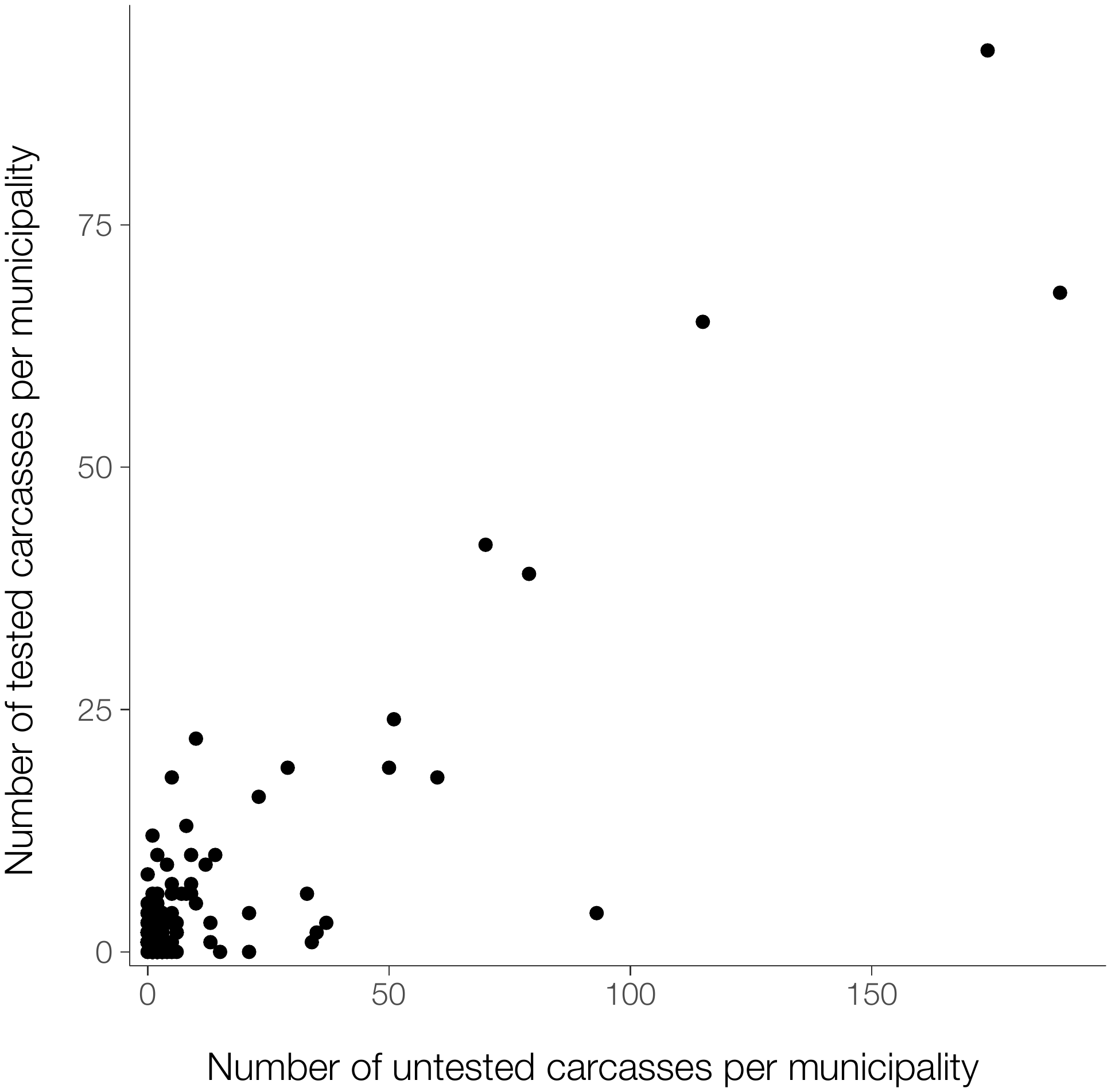

**Figure S2.** Number of carcasses tested and untested for yellow fever virus per municipality.

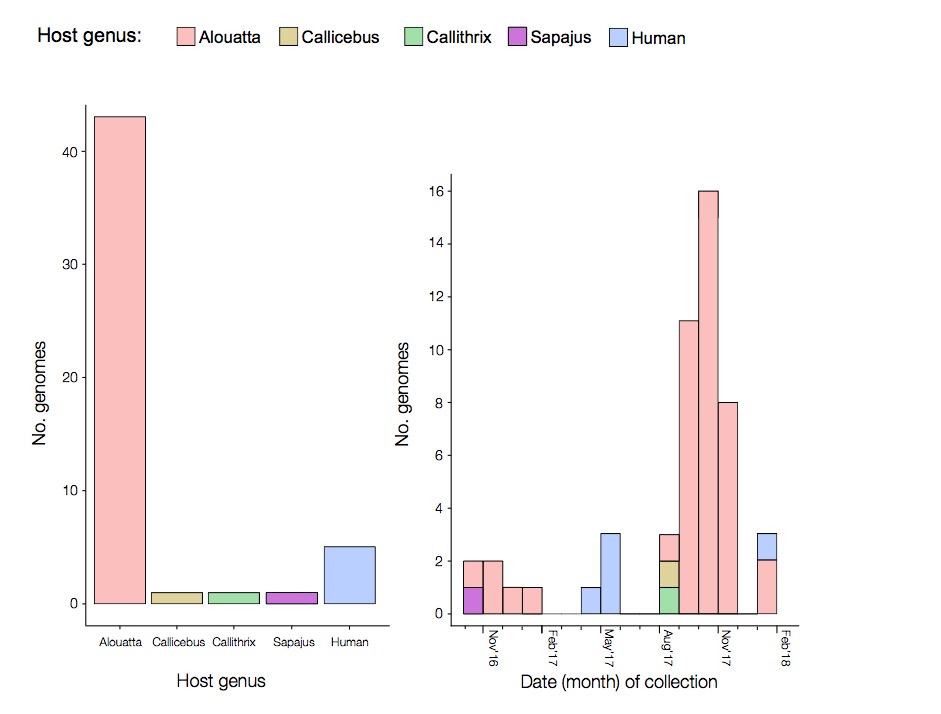

**Figure S3.** Date of sample collection and number of genomes generated from each host genus.

**­­­­**
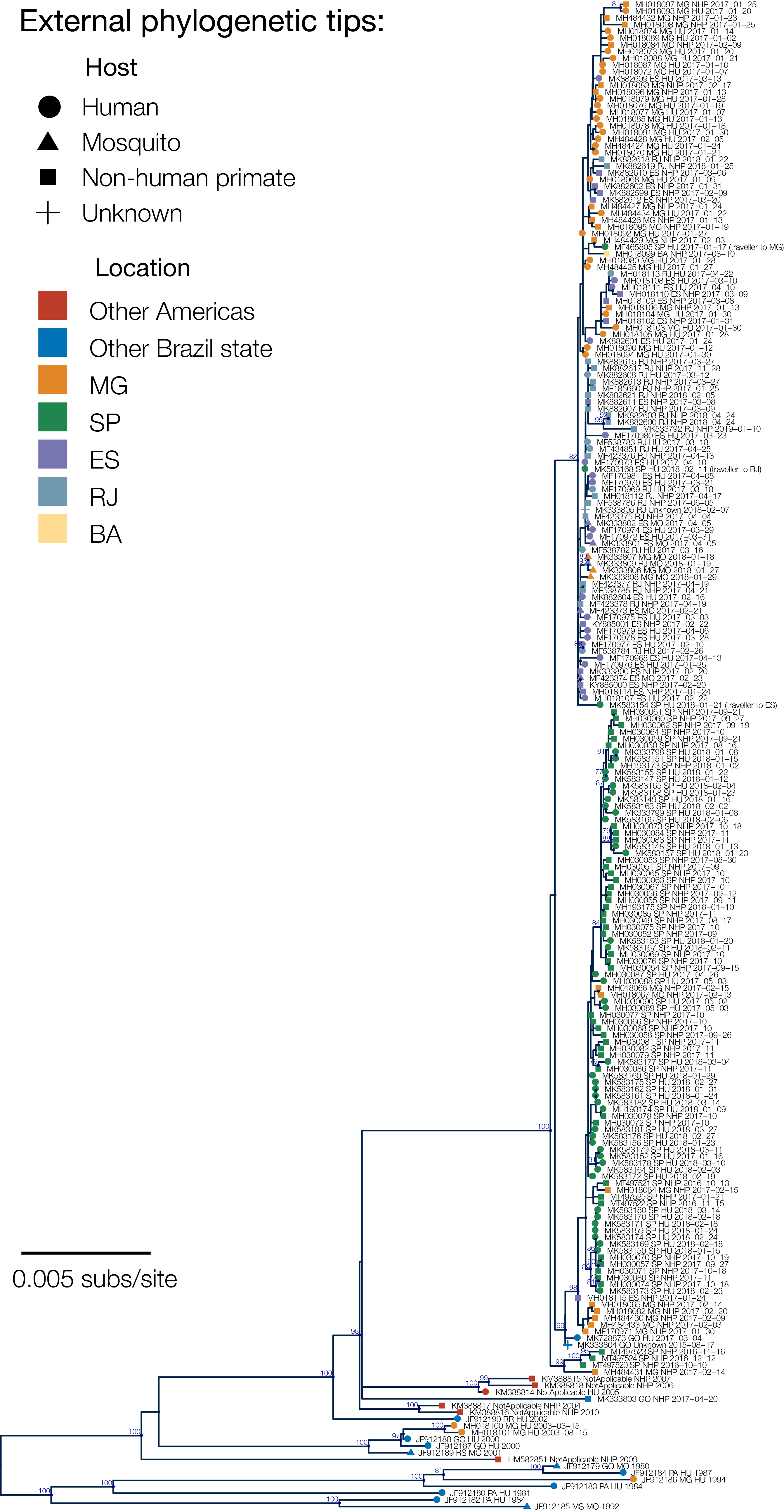

**Figure S4.** Maximum likelihood phylogeny of new generated genomes sequences and all publicly available SA1 YFV sequences from Brazil (n=224).

**Supplementary Tables**

**Table S1.** Details of the YFV genome generated in this study. All sequences are from São Paulo.

| **Accession number** | **Sequence ID** | **Municipality** | **Host** | **Collection date** | **CT** |
| --- | --- | --- | --- | --- | --- |
| MT497520 | CP224 | Potirendaba | Sapajus | 10 Oct 2016 | 22 |
| MT497521 | CP159 | Pindorama | Alouatta | 13 Oct 2016 | 18 |
| MT497522 | CP182 | Jaboticabal | Alouatta | 15 Nov 2016 | 15 |
| MT497523 | CP227 | Catanduva | Alouatta | 16 Nov 2016 | 12 |
| MT497524 | CP164 | Catigua | Alouatta | 12 Dec 2016 | 14 |
| MT497525 | CP179 | Ribeirao Preto | Alouatta | 21 Jan 2017 | 16 |
| MH030087 | Y39 | Americana | Human | 26 April 2017 | 32 |
| MH030090 | Y42 | Sao Joao Boa Vista | Human | 02 May 2017 | 35 |
| MH030088 | Y40 | Tuiuti | Human | 03 May 2017 | 37 |
| MH030089 | Y41 | Sao Joao Boa Vista | Human | 03 May 2017 | 33 |
| MH030050 | Y2 | Louveira | Callicebus | 16 August 2017 | 11 |
| MH030049 | Y1 | Vinhedo | Callithrix | 17 August 2017 | 12 |
| MH030053 | Y5 | Itatiba | Alouatta | 30 August 2017 | 10 |
| MH030051 | Y3 | Louveira | Alouatta | Sept 17 | 9 |
| MH030052 | Y4 | Louveira | Alouatta | Sept 17 | 15 |
| MH030055 | Y7 | Jundiai | Alouatta | 11 Sept 2017 | 11 |
| MH030056 | Y8 | Jundiai | Alouatta | 12 Sept 2017 | 9 |
| MH030054 | Y6 | Jundiai | Alouatta | 15 Sept 2017 | 12 |
| MH030062 | Y14 | Itatiba | Alouatta | 19 Sept 2017 | 12 |
| MH030059 | Y11 | Itatiba | Alouatta | 21 Sept 2017 | 12 |
| MH030061 | Y13 | Itatiba | Alouatta | 21 Sept 2017 | 13 |
| MH030058 | Y10 | Itatiba | Alouatta | 26 Sept 2017 | 10 |
| MH030057 | Y9 | Braganca Paulista | Alouatta | 27 Sept 2017 | 16 |
| MH030060 | Y12 | Itatiba | Alouatta | 27 Sept 2017 | 12 |
| MH030063 | Y15 | Itatiba | Alouatta | Oct 17 | 11 |
| MH030064 | Y16 | Jundiai | Alouatta | Oct 17 | 10 |
| MH030065 | Y17 | Jundiai | Alouatta | Oct 17 | 12 |
| MH030066 | Y18 | Jundiai | Alouatta | Oct 17 | 11 |
| MH030067 | Y19 | Jundiai | Alouatta | Oct 17 | 24 |
| MH030068 | Y20 | Jundiai | Alouatta | Oct 17 | 12 |
| MH030069 | Y21 | SaoPaulo | Alouatta | Oct 17 | 16 |
| MH030072 | Y24 | Jarinu | Alouatta | Oct 17 | 10 |
| MH030075 | Y27 | Jarinu | Alouatta | Oct 17 | 19 |
| MH030076 | Y28 | Jarinu | Alouatta | Oct 17 | 17 |
| MH030077 | Y29 | Jarinu | Alouatta | Oct 17 | 16 |
| MH030078 | Y30 | Morungaba | Alouatta | Oct 17 | 12 |
| MH030071 | Y23 | Campo Limpo Paulista | Alouatta | 18 Oct 2017 | 21 |
| MH030073 | Y25 | Mairipora | Alouatta | 18 Oct 2017 | 24 |
| MH030074 | Y26 | Jarinu | Alouatta | 18 Oct 2017 | 15 |
| MH030070 | Y22 | Campo Limpo Paulista | Alouatta | 19 Oct 2017 | 14 |
| MH030079 | Y31 | Nazare Paulista | Alouatta | Nov 17 | 15 |
| MH030080 | Y32 | Nazare Paulista | Alouatta | Nov 17 | 14 |
| MH030081 | Y33 | Campo Limpo | Alouatta | Nov 17 | 25 |
| MH030082 | Y34 | Campo Limpo | Alouatta | Nov 17 | 16 |
| MH030083 | Y35 | Campo Limpo | Alouatta | Nov 17 | 15 |
| MH030084 | Y36 | Piracaia | Alouatta | Nov 17 | 13 |
| MH030085 | Y37 | Piracaia | Alouatta | Nov 17 | 14 |
| MH030086 | Y38 | Piracaia | Alouatta | Nov 17 | 15 |
| MH193173 | SA129 | Guarulhos | Alouatta | 02 Jan 2018 |  |
| MH193174 | SA130 | Mairipora | Human | 09 Jan 2018 |  |
| MH193175 | SA131 | Sao Paulo | Alouatta | 10 Jan 2018 |  |

**Table S2. Non-human primate yellow fever virus genome sequences from São Paulo, by host genus.** Detailed information for each sequenced isolate can be found in **Table S1**.

| **Host genus** | **Confirmed NHP cases*** | | | **Genomes generated** | | | **qPCR-Ct values** | | | **Sequence coverage** | | |
| --- | --- | --- | --- | --- | --- | --- | --- | --- | --- | --- | --- | --- |
|  | Total | % of total** | | Total | % of total | | Median | | Range | Median | | Range |
| Alouatta | 403 | | 88 | 43 | | 84 | 14 | 9-25 | | 99.3 | 86.1-99.4 | |
| Callicebus | 9 | | 2 | 1 | | 2 | 11 | n.a. | | 99.3 | n.a. | |
| Callithrix | 35 | | 8 | 1 | | 2 | 12 | n.a. | | 99.2 | n.a. | |
| Cebidae | 9 | | 22 | 0 | | 0 | n.a. | n.a. | | n.a. | n.a. | |
| Sapajus | 3 | | 0.7 | 1 | | 2 | 22 | n.a. | | 96.3 | n.a. | |
| Human | - | | - | 5 | | 10 | 34 | 32-37 | | 96.1 | 80.8-99.3 | |

*Cases reported to Instituto Adolfo Lutz, São Paulo, from week 29 of 2016 through week 4 of 2018. n.a.=not applicable.

**For which genera was known

**Table S3: Mean branch dispersal velocity estimates.**

| **Dataset (number of sequences)** | **Mean branch dispersal velocity (km/day, [95% HPD]) based on phylogenetic branches in defined phases** | | | |
| --- | --- | --- | --- | --- |
|  | **All** | **Phase 1: root up to 2017-02-01** | **Phase 2: 2017-02-01 to 2017-07-01** | **Phase 3: 2017-07-01 onwards** |
| **Full (99)** | 0.95 (0.53-2.42) | 1.87 (0.64-5.95) | 1.05 (0.23-2.89) | 0.80 (0.44-1.85) |
| **Geographically restricted (95)** | 0.83 (0.50-1.53) | 0.86 (0.40-1.94) | 0.93 (0.27-2.58) | 0.87 (0.46-1.79) |
